# New insights into the systematics of the Afrotropical *Amblyomma marmoreum* complex (Acari, Ixodidae) and a novel *Rickettsia africae* strain using morphological and metagenomic approaches

**DOI:** 10.1101/2023.08.18.553479

**Authors:** Andrea P. Cotes-Perdomo, Alberto Sánchez-Vialas, Richard Thomas, Andrew Jenkins, Juan E. Uribe

**Affiliations:** Faculty of Technology, Natural Sciences and Maritime Sciences, Department of Natural Sciences and Environmental Health, University of South-Eastern, Norway; Department of Biodiversity and Evolutionary Biology, Museo Nacional de Ciencias Naturales (MNCN-CSIC), Madrid, Spain; Facultad de Ciencias Veterinarias, Departamento de Ciencia Animal, Universidad de Concepción, Chillán, Chile

**Keywords:** Illumina Novaseq6000, Mitochondrial genome, African tick-bite fever, Spotted fever group, *Rickettsiales* bacteria, Tick metagenomics

## Abstract

The *Amblyomma marmoreum* complex includes some Afrotropical species, such as *Amblyomma sparsum*, a three-host tick that parasitizes reptiles, birds, and mammals, and is a recognized vector of *Ehrlichia ruminatum*. However, the lack of morphological, genetic and ecological data on *A. sparsum* has caused considerable confusion in its identification among *A. marmoreum* complex members. In this study, we used microscopy and metagenomic approaches to analyze *A*. sparsum ticks collected from a puff adder snake (*Bitis arietans*) in southwest Senegal (an endemic rickettsioses area) in order to supplement previous morphological descriptions, provide novel genomic data for the *A. marmoreum* complex, and search for some associated spotted fever agent. Based on stereoscope and scanning electron microscopy (SEM) morphological evaluations, we provided high-quality images and new insights about punctation and enameling in the male of *A. sparsum* to facilitate identification for future studies. Additionally, the metagenomic approach allowed us assembly the complete mitochondrial genome of *A. sparsum*, as well as the nearly entire chromosome and complete plasmid sequences of a novel *Rickettsia africae* strain. Phylogenomic analyses showed a close relationship between *A. sparsum* and *A. nuttalli* for the first time and confirmed the position of *A. sparsum* within the *A. marmoreum* complex. Our results provide new insights into the systematic of *A. sparsum* and *A. marmoreum* complex, as well as the genetic diversity of *R. africae* in Afrotropical region. Future studies should consider the possibility that *A. sparsum* may be a competent vector for *R. africae*.

## 1. Introduction

Ticks (Acari: Ixodida) are hematophagous ectoparasites that infest vertebrates; they are important carriers of infectious diseases and can transmit a plethora of pathogenic bacteria, viruses, and protistan parasites (Baneth, 2014). Lyme, anaplasmosis, rickettsiosis, ehrlichiosis, and babesiosis, are among some of the tick-borne diseases, which have significant economic and health impacts (Jongejan and Uilenberg, 2004). The order Ixodida includes some 953 species (Guglielmone et al., 2020) clustered into the Ixodidae (758 spp.), the Argasidae (212 spp.) and the monotypic Nutalliellidae (ITIS, 2023, Guglielmone et al., 2020).

*Amblyomma* is the third-largest genus within Ixodidae, with 136 species, widely distributed in America, Sub-Saharan Africa, Asia and Australasia (Guglielmone et al., 2020). Multiple species complexes have been recognized within *Amblyomma*, including the *A. cajennense* and *A. maculatum* complexes in America, the *A. testudinarium* complex in Asia, and the *A. marmoreum* complex in Africa (Guglielmone et al., 2017). Morphological identification within these complexes has proved challenging, and despite several molecular and morphological studies in recent years (Beati et al., 2013; Nava et al., 2014; Lado et al., 2018; Mohamed et al., 2022) much remains to be done particularly in respect of the species complexes in Africa and Asia.

The *Amblyomma marmoreum* complex comprises five African species: *A. marmoreum* sensu stricto, *A. sparsum, A. falsomarmoreum, A. nuttalli, and A. paulopunctatum. Amblyomma sparsum* occurs in parts of central and eastern Africa, overlapping with *A. marmoreum* in northern Zimbabwe (Walker and Olwage, 1987). As a three-host tick, adults primarily parasitize large reptiles and occasionally birds and large mammals, while the immature stages favour small mammals (Petney et al., 1987; Guglielmone et al., 2014). Unfortunately, the lack of morphological, genetic, and ecological data on the *Amblyomma marmoreum* complex members has caused considerable confusion in their identification for decades (Theiler and Salisbury, 1959; Guglielmone et al., 2017), and their evolutionary relationships, distribution, vector potential and hosts remain incompletely delineated.

Importantly, some species of the *A. marmoreum* complex, along with their reptilian and mammalian hosts, have the potential to be involved in the infectious cycles of pathogens, such as *Rickettsia* spp. (*Rickettsiales*: *Rickettsiaceae*) (Mendoza-Roldan et al., 2021), which are members of the *Alphaproteobacteria* which cause acute febrile illnesses with potentially severe or fatal outcomes, such as spotted fever group (SFG) and typhus (Raoult et al., 2005).

In this study, we analyze male *Amblyomma sparsum* ticks collected from a puff adder snake (*Bitis arietans*) in southwest Senegal, where tick-borne rickettsioses are emerging diseases in rural areas (Mediannikov et al., 2010). In order to supplement the hitherto limited morphological and molecular information on this species we performed detailed morphological comparisons with previous descriptions and used a metagenomic approach to determine the mitochondrial genome sequence. This allowed us to review the phylogeny of the *Amblyomma marmoreum* complex. In addition, we were able to determine the genomic and plasmid sequence of an associated *Rickettsia* species, which was found to be a novel strain of *R. africae*.

## 2. Material and methods

### 2.1. Sample collection, morphological identification, and DNA extraction

Four male ticks were collected from a puff adder in southwest Senegal (13.623339, -16.430217), (Figure 1) on August the 26^th^, 2015, and stored in sterile tubes with 96% ethanol. The ticks were identified as *Amblyomma sparsum* males on the basis of morphological characteristics described by Theiler and Salisbury (1959) for the *A. marmoreum* complex.

**Figure 1.**
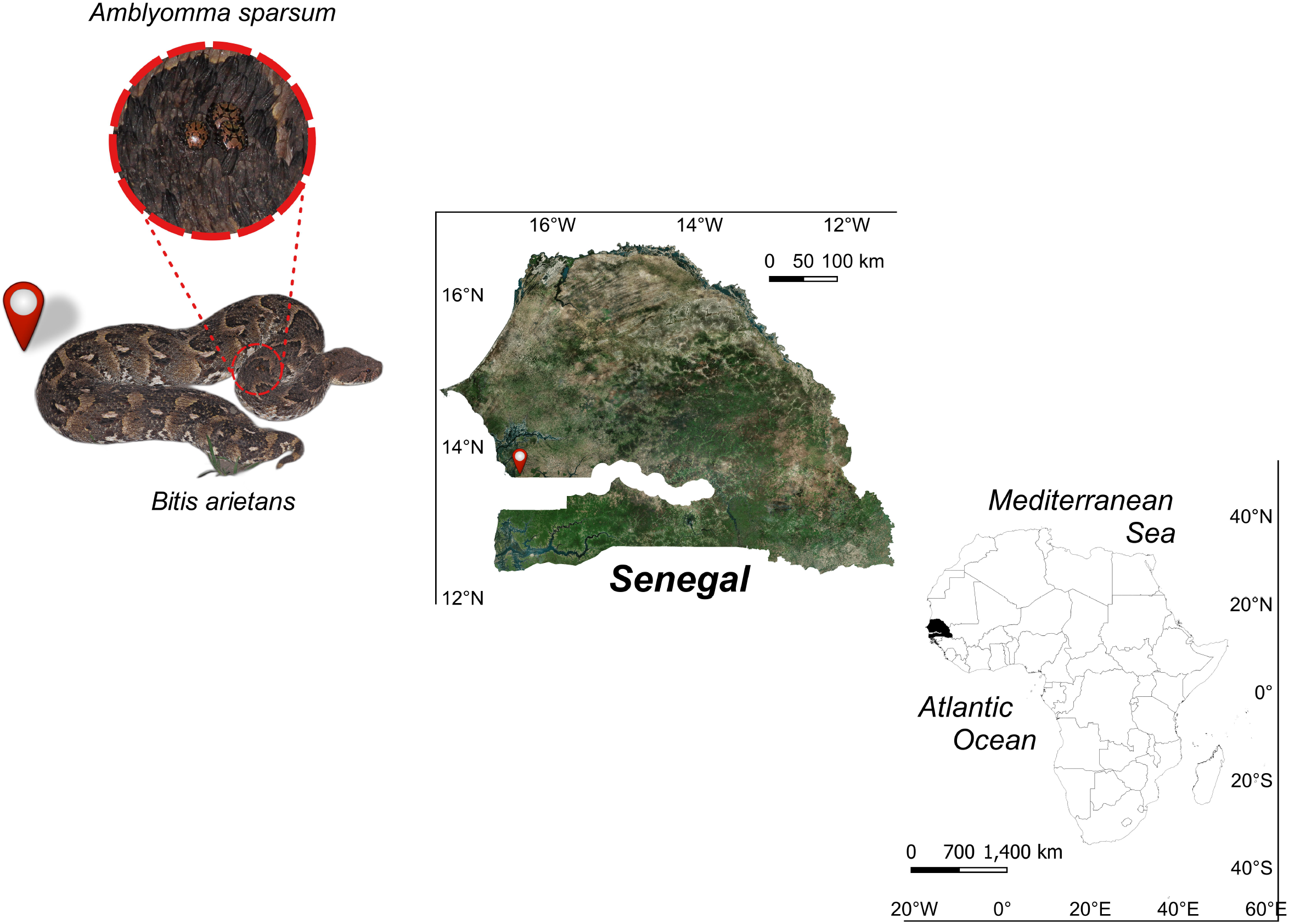
From left to right are shown *i: Bitis arietans* snake hosting the specimens of *Amblyomma sparsum, ii: s*ampling location in Senegal, and *iii:* its African context.

Two specimens were subjected to a detailed morphological assessment using a Canon 700D camera fitted with a macro lens and two external flashes coupled with a stereoscope and a Scanning Electron Microscope (SEM) at the Museo Nacional de Ciencias Naturales (MNCN-CSIC), Madrid, Spain.

Genomic DNA was extracted from one specimen using the Dneasy Blood & Tissue kit (Qiagen, Valencia, CA, USA), following the manufacturer’s instructions but with the initial proteinase K lysis step extended to 16 hours at 56°C with continuous agitation.

### 2.2. Massive sequencing and genomic data analysis

The DNA was used to construct a Truseq Nano DNA Library, which was sequenced using the Illumina platform (Novaseq6000 150 pair-end/10Gb). The raw sequencing data were subjected to quality control using FastQC (Andrews, 2010). Adapter sequences were trimmed off, redundant and/or unpaired sequences were removed, and low-quality reads were discarded using Trimmomatic (Bolger et al., 2014). Cleaned fastq files were then assembled *de novo* using SPAdes (Prjibelski et al., 2020) with default parameters.

To extract the mitogenome of the tick and any SFG rickettsial species present, we developed two custom genome-level databases using “makeblastdb” tool of BLAST (Altschul, 1997). The first database included all available mitogenomes for *Amblyomma* ticks. The second contained the genomic sequences of SFG *Rickettsia* species (Abubakar et al., 2019). The assembled sequences were faced against the custom databases using “blastn” command tool of BLAST. Then, the “subseq” command of seqtk (Li, 2012) tool was used to identify reads that corresponded to mitochondrial and *Rickettsia* genomes respectively. Subsequently, the genomic information from each search was mapped with Geneious^®^ to visually check assembly and mean depth of coverage.

Mitochondrial sequences were annotated using MITOS (Bernt et al., 2013) and the open reading frames (ORFs) were manually checked. The boundaries of ribosomal genes were determined following Boore et al., (2005). The amino acids composition, CG percentage, genetic distance comparison, and relative synonymous codon usage (RSCU) were analyzed for a matrix of 17 mitochondrial genomes of *Amblyomma* species using MEGA 11 (Tamura et al., 2021). Raw read sequence data have been deposited in the NCBI Sequence Read Archive (SRA: SRR24067184) database (BioProject: PRJNA951762).

### 2.3. Phylogenomic analysis

For phylogenomic analysis, the *Amblyomma sparsum* mitogenome determined in this study was aligned with available mitogenome data of *Amblyomma* species in GenBank, using other genera of Ixodida as outgroups (Supplementary Material 1). In order to optimize comparison of protein-coding genes, homologous ORFs were aligned using the Translator X server (Abascal et al., 2010) with nucleotide codification based on the invertebrate mitochondrial genetic code. The alignments and subsequent alignment curation were performed using MAFFT (Katoh et al., 2005) and GBlocks (Castresana, 2000), respectively, with default parameters. Subsequently, a concatenated matrix of the 13 protein-coding genes and two ribosomal genes (at nucleotide level) was created and analyzed using Bayesian inference (BI: Rannala and Yang, 1996; Yang and Rannala, 1997) and Maximum Likelihood (ML: Felsenstein, 1981) methods.

BI analysis was run with PhyloBayes MPI v. 1.5a (Lartillot et al., 2013) using the GTR-CAT heterogeneity model on two independent Markov chain Monte Carlo, sampling at every cycle until convergence, which was checked with Tracer v. 1.7.1 (Rambaut et al., 2018). ML trees were constructed using IQ-TREE v. 1.6.1 (Nguyen et al., 2014), employing a combination of rapid hill-climbing and stochastic perturbation methods. The robustness of the inferred trees was assessed using 1,000 pseudoreplicates of ultrafast bootstrap (UFBoot). The best-fit mixture models (mtZOA + F + C50 + R5) were selected for ML analyses based on Bayesian information criteria (Schwarz, 1978) in ModelFinder (Kalyaanamoorthy et al., 2017) with the following commands: “-m TESTONLY” to choose the best empirical evolutionary model (EM) and “-mset EM + C10, EM + C20, EM + C30, EM + C40, EM + C50, EM + C60”.

SFG *Rickettsia* sequence information extracted by the BLAST search was assessed for completeness using an *Alphaprotobacteria* database (alphaproteobacteria_odb10; Creation date: 2020-03-06, number of genomes: 744, number of BUSCOs: 432) using BUSCO (Manni et al., 2021). Finally, the *Rickettsia* genomic information was annotated using Prokka version v. 1.14.6 (Seemann, 2014) with default parameters, using the chromosome and plasmid sequences of *Rickettsia africae* ESF-5 (CP001612 and CP001613, respectively) as templates.

The resulting *Rickettsia* genome, together with other genomes of *Rickettsia* species available in GenBank and belonging to the same lineage (El Karkouri et al., 2022), were used to construct a phylogenetic tree (Supplementary Material 2). The single-copy genes of each genome were determined using BUSCO. These were then assembled into FASTA files (one FASTA file per species) and orthologous groups were inferred with OrthoFinder 2.5.4 (Emms and Kelly, 2019). The orthologs were concatenated into a single matrix and an ML tree was inferred as described above. Four *Rickettsia* species (*R. agasii, R. fourneri, R. gravesii and R. honei)* were excluded from the analysis as they contained potential gene duplications detected in BUSCO (see Supplementary Material 3).

## 3. RESULTS

### 3.1. Morphological identification and characterization

The four ticks were identified morphologically as male *Amblyomma sparsum* based on the description in Theiler & Salisbury (1959). Main differences between *A. sparsum* and other species in the *A. marmoreum* complex are: (i) the marginal groove, which is not as deep as in the other four species and is absent in the anterior portion; and (ii) the enamel and stripe patterns, which differ principally in the festoons (Figure 2).

**Figure 2.**
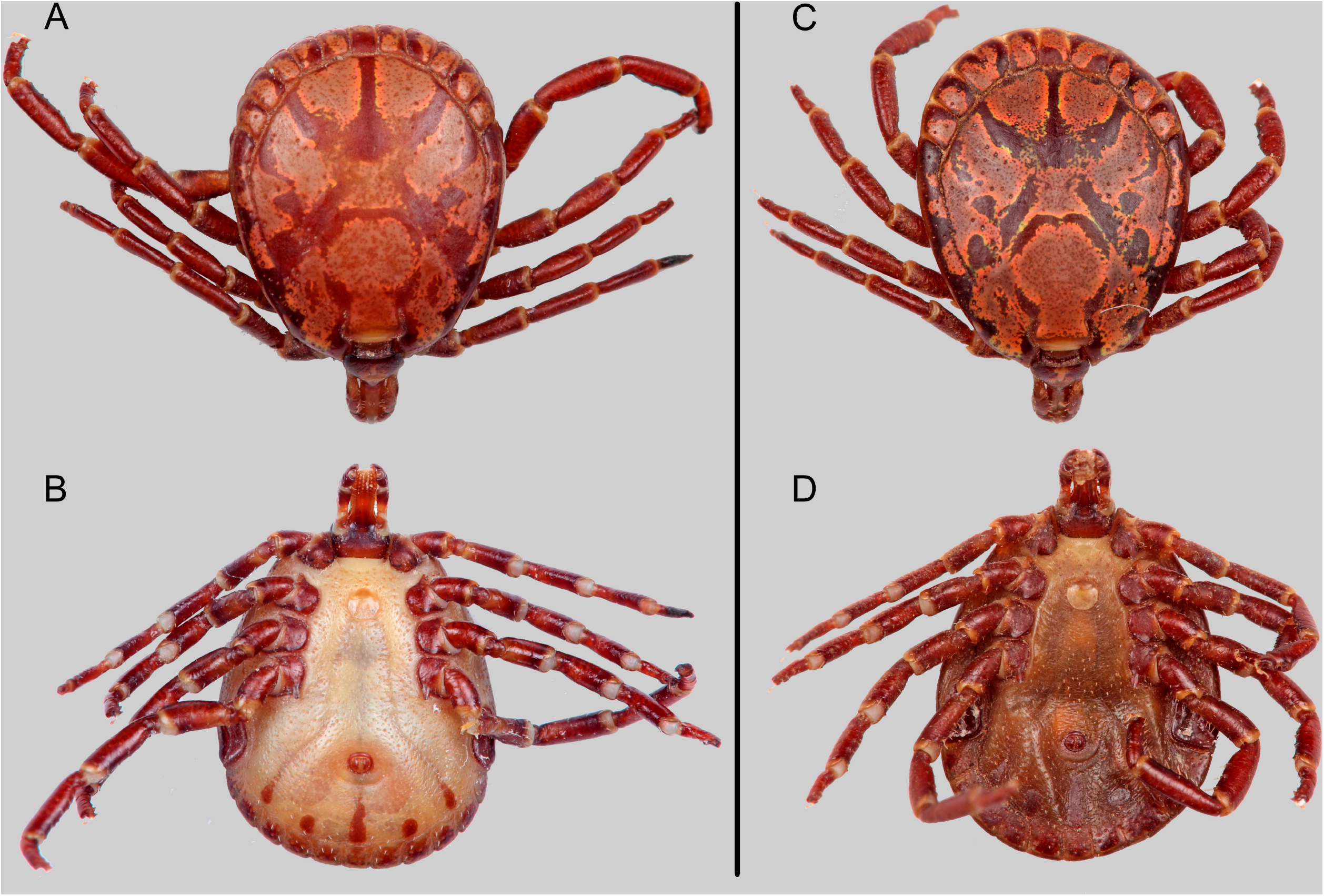
Males of *Amblyomma sparsum* A. and B. dorsal and ventral view of specimen M1; C and D. dorsal and ventral view of M2.

Figure 2 shows the ornate, ovoid conscutum, which displays a similar enameling pattern and coloration in both specimens. Both specimens have a copper-green iridescence, although this is barely visible in specimen M1 (Figure 2: A and C). Lateral grooves are absent. Cervical grooves are short, moderately deep, and have a semi-concave line shape (Figure 3: A). The eyes of both individuals are yellow to light brown and appear flush with the surface of the conscutum, although a slight relief is revealed in Figure 3: A.

**Figure 3.**
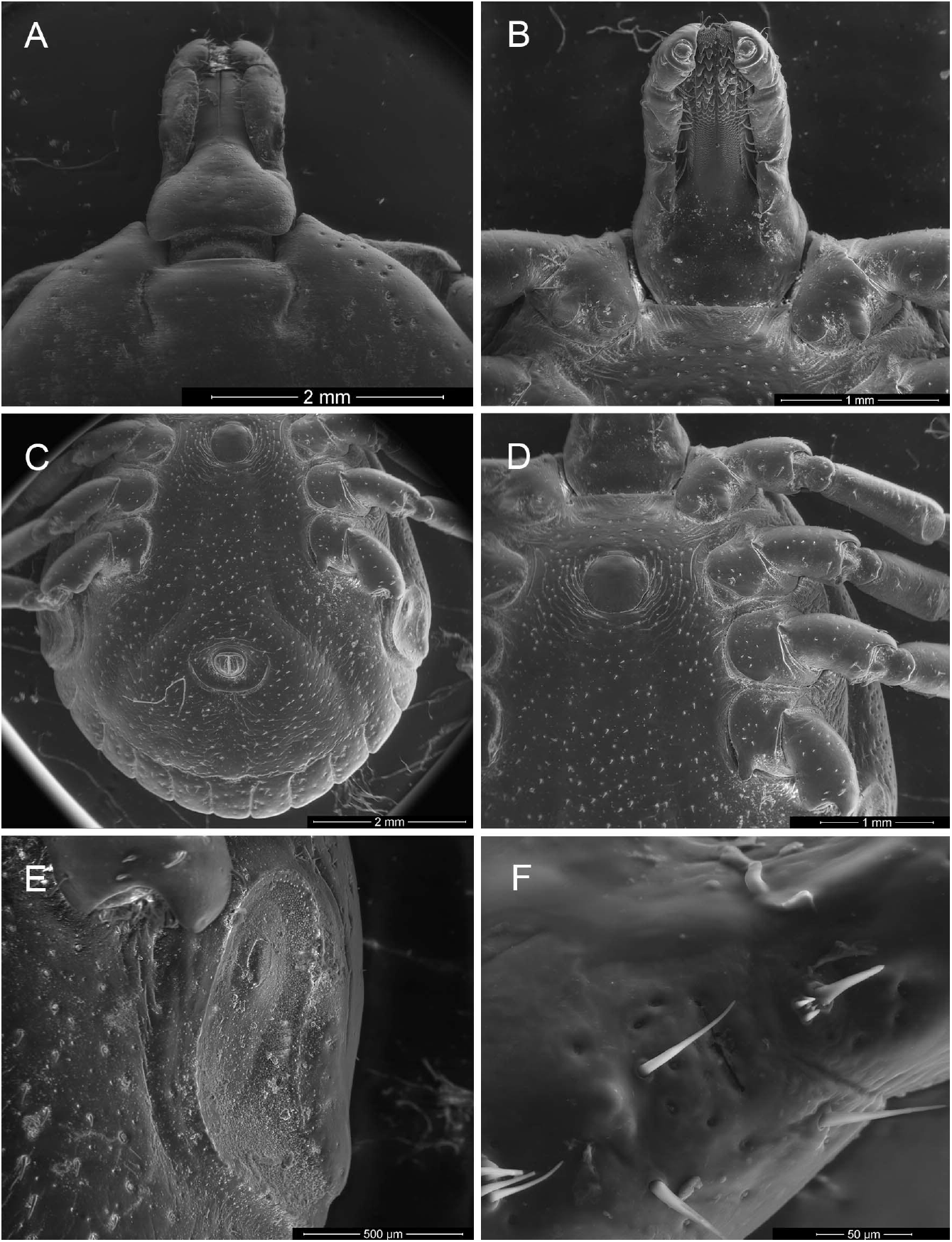
A. and B. dorsal and ventral view of the capitulum; C. anal pore and festoons; D. coxae and genital aperture; E. spiracular plate; and F. Haller’s organ.

Both specimens display a marginal groove that shares similar characteristics with the type specimen described by Neumann (1899) (see Theiler and Salisbury, 1959). In this regard, the anterior portion of the groove consists of a row of punctations that gradually become discontinuous. The festoons are pronounced and have a longer-than-broad shape. The specimen M2 (Figure 2: C and D) shows a fusion of two festoons, resulting in a reduced distinctiveness both dorsally and ventrally. However, both specimens display a similar enameling pattern in their festoons. Spiracular plates are comma-shaped, and their caudal processes are slightly longer than the adjacent festoons (Figure 3: E). Ventrally, there are five plaques; the central plaque and the outer two are lanceolate, with the central plaque being longer. The intermediate plaques are shorter and almost circular (Figure 2). The genital aperture is located between the coxae II and has a circular shape (Figure 3: D).

The fovea, which is not readily visible, is centrally located between the falciform stripes and near the tip of the posteromedian stripe (Figure 2). Both specimens present numerous, large punctations over the entire conscutum, particularly in the posterior and lateral parts, including the shoulders. A small tubercule is visible in most of the punctations. The Haller’s organs present a transverse aperture (Figure 3: F). The basis of the capituli has small, centered punctations and is roughly as broad as long. A heart-shaped enameling pattern is present on the basis of capituli in both specimens, with the thinner end reaching the posterior edge. The palps present a pale enamel on the dorsal side. Short, wide, rounded subcollars are present. The two specimens presented a 3/3 dentition in most of the anterior hypostome, but 2 or 3 rows at the basis presented a 3.5/3.5 dentition, the corona is filled with fine denticles (Figure 3: B).

In our specimens, the spots between the antero-accessory stripes and the postero-accessory stripes meet the antero-accessory stripes, and at the same time, the antero-accessory stripe touches the lateral groove, reaching the third lateral spot. The falciform stripe is unbroken in the specimen M1 (Figure 2: A) but broken in the mid-line of the specimen M2 (Figure 2:C). Although the four lateral spots are present in both individuals, they are not equally delimited and are more clearly defined in specimen M1 (Figure 2: A). The space between the cervical and falciform stripes is short and almost closed in specimen M1 (Figure 2: A). In addition, both the color and SEM photographs show that the spurs of Coxae II and III are more pointed and slightly more elongated than in the original description. The spur in Coxa IV is rounded, with a constant width, and thus less pointed than the original description (Figure 3: D).

### 3.2. General features of the *Amblyomma sparsum* mitochondrial genome

The mitochondrial sequence (GenBank record OQ842962) corresponds to a double-stranded circular molecule with a length of 14,679 bp. The base composition 21.1% GC. It contains 37 genes, coding for 13 proteins, 2 rRNAs and 22 tRNAs, plus two control regions of respectively 306 bp, between the 12S rRNA and tRNA^Ile^, and 270 bp, between tRNA^Leu^ and tRNA^Cys^. Twenty-four genes overlap: ATP8 and ATP6, ND4L and ND4, ND1 and tRNA^Glu^ by seven bases; NAD2 and tRNA^Trp^, tRNA^Trp^ and tRNA^Tyr^ by two bases; ND3 and tRNA^Ala^, ND4 and tRNA^His^, tRNA^Lys^ and tRNA^Asp^, tRNA^Arg^ and tRNA^Asn^, tRNA^Asn^ and tRNA^Ser^, CYTB and tRNA^Ser^, tRNA^Ser^ and tRNA^Leu^ by one base. Two genes (ND4 and CYTB) have TAG as a stop codon, ten (ND5, COI, ND2, ND1, COII, ATP6, ND6, ND3, ND4L, and ATP8) have TAA, while the COIII gene has a truncated the stop codon (T––) that needs polyadenylation to function as a TAA stop codon (See Supplementary Material 4).

### 3.3. Molecular identification

Comparison of our mitochondrial sequence with sequences available on GENBANK showed 89.1% and 87.1% identity with the two entire mitogenomes of *A. marmoreum*, KY457515 and KY457515, respectively (Mans et al., 2019), and 95.6% identity with *A. nuttalli* (OL741736; Kelava et al., 2023). Only two DNA sequences attributed to *A. sparsum* were found; an 18S rRNA (MF102090; Beati and Klompen, 2019) showing 100% identity with our sequence, and a 12S rRNA (AF150047; Beati and Keirans, 2001) showing only 88.2% identity. A BLAST search of this sequence showed a high similarity with the 12S rRNA partial genes of *A. variegatum* available in the database. It was collected in an area where both *A. sparsum* and *A. variegatum* are known to occur.

### 3.4. Phylogenetic analysis

ML and BI analyses gave slightly different tree topologies, the principal differences being in the positions of the species of the Indo-Pacific *A. testudinarium* group and African *Amblyomma* species. In both trees, *A. sparsum* groups with *A. nuttalli* as the sister group of *A. marmoreum*. There is good support (88 of Bootstrap Percentage –BP–/0.97 Posterior Probability –PP–) for inclusion of *A. hebraeum* and *A. tholloni* (Figure 4, left tree) within the group. However, the phylogenetic position of the *A. testudinarium* complex relative to *A. hebraeum* and *A. tholloni* remains unresolved (Figure 4, left tree). *A. ovale* and *A. maculatum* are deeply-branching taxa, but their exact position relative to each other and the *A marmoreum* and *A. testudinarium* groups is not resolved. *A. triguttatum* consistently appears as a basal taxon in both trees. The position of the outgroup branch is stable, indicating that *Amblyomma* is monophyletic.

**Figure 4.**
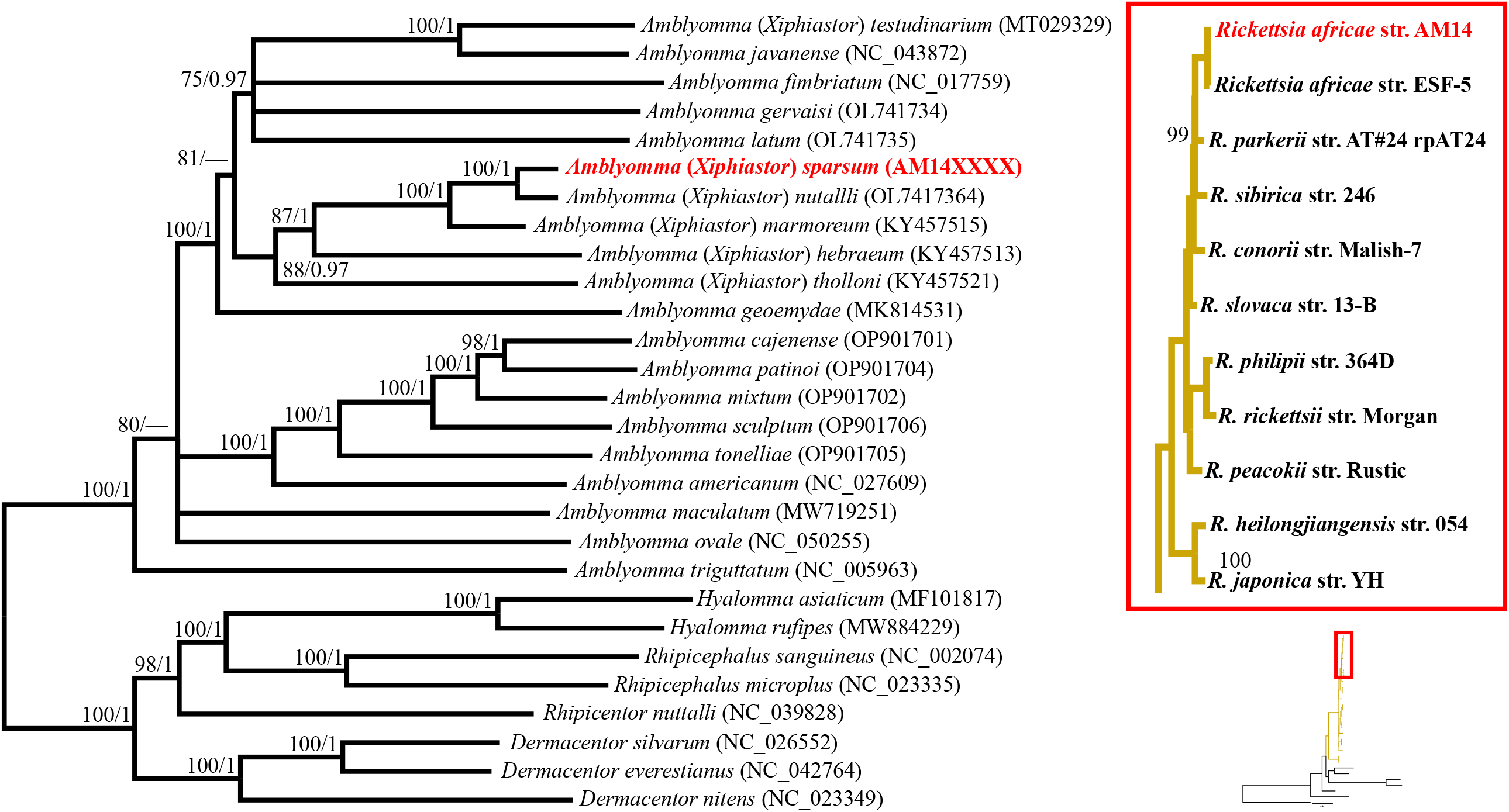
Phylogenomic relationships of the *Amblyomma* genus (left) and phylogenomic relationships of the *Rickettsia* species (right), based on mitochondrial and genomic data. The left tree is based on 13 mitochondrial protein-coding and rRNA genes at the nucleotide level. The numbers at nodes represent statistical support for maximum likelihood (ML) and Bayesian inference analyses, respectively. Nodes with support values below 75 for bootstrap and 0.95 for posterior probability are collapsed. The right tree displays the ML phylogenomic relationships of *Rickettsia* species based on 221 homologous protein-coding genes. The red inset highlights the relationships recovered for *Rickettsia africae*-related species. Nodes without bootstrap values in the ML tree were recovered with maximum support (100).

#### Associated Spotted Fever *Rickettsia* genome and its phylogenetic position

Analysis of *Rickettsia* sequences extracted from the library identified the complete genome of a *Rickettsia africae* strain which we term Senegal-AM14. The genome has a total length of 1,281,475 bp, a GC content of 32.9% and 98.7% identity with an existing *R. africae* genome sequence (strain ESF-5). A total of 1,470 potential genes were found, including 1,434 protein-coding genes, 33 tRNAs, and 3 rRNAs. Additionally, a circular plasmid of 8,801 bp with eight protein-coding genes was found and shown to be highly homologous to the plasmid of ESF-5, although it is shorter and contains four fewer genes. The graphic summary of BUSCO completeness of all *Rickettsia* analyzed against alphaproteobacterial_odb10 is shown in Supplementary Material 4. The current sequence is 74.5% complete. Phylogenetic analysis indicates that *R. africae* AM14 and ESF5 group together as a sister taxon of *R. parkerii* (inset of Figure 4).

## 4. DISCUSSION

Although *Amblyomma sparsum* is commonly found on reptiles, mammalian hosts including cattle are also frequent hosts (Theiler and Salisbury, 1959; Petney et al., 1987) making it a potential vector for pathogens that are important for public health such as *Rickettsia africae*, and for livestock, such as *Ehrlichia ruminantium* (Walker and Olwage, 1987; Burridge et al., 2000). Thus, its phylogeny, identification and phylogeography may be of public health and economic interest.

The puff adder (*B. arietans*) is a common host for *Amblyomma sparsum* (Theiler and Salisbury, 1959). It has a pan-African distribution, mostly in sub-Saharan regions, but with established populations in the western Sahara and the southern part of the Arabian Peninsula (Barlow et al., 2013), which parallels the recorded distribution of *A. sparsum* and its apparent ecological preferences for arid and semi-arid environments (Petney et al., 1987; Guglielmone et al., 2014).

Studies of the genetic structure of mammalian and reptile species with transcontinental distributions have revealed the presence of climatically determined barriers and refugia acting differently on different groups (Barlow et al., 2013; Smitz et al., 2013). Studying the genetic structure of tick populations could cast further light on this issue and the potential impact of future climate scenarios. The establishment of robust phylogenetic frameworks is a prerequisite for such studies (Beati and Klompen, 2019).

Genomic information from ticks and their associated microorganisms is a key to the understanding of their biology, ecological interactions, detection of potential pathogens, and thus the development of measures that allow assessing and predicting the zoonotic risks in the climate change scenarios (Nuttall, 2022).

### Morphology and molecular identification

The four ticks investigated had a morphology agreeing with the original description of *A. sparsum*, although some variation was present, especially in the enameling pattern. It is important to map such variation as ornamentation patterns are assigned high taxonomical importance in other species complexes, such as the *A. cajennense* and *Rhipicephalus sanguineus* complexes along with genital aperture shape and adjacent structures, color, spiracular plate shape and length, and adanal plates (Nava et al., 2014, Feldman-Muhsam, 1952; Dantas-Torres et al., 2013).

Theiler and Salisbury (1959) provide a valuable and comprehensive description of the species within the *A. marmoreum* complex, including variations related to methods of preservation and observation. Despite this, diagnosis remains a challenge, even leading to confusion with ticks outside the complex (Beati and Keirans, 2001), and this may explain why a 12S rRNA sequence that the latter authors attribute to *A. sparsum* resembles *A variegatum* sequences rather than that which we report. Additional imaging techniques such as electron microscopy and high-quality stereoscopic pictures may help to delineate additional characteristics such as punctation and enameling, which may facilitate identification and future studies.

*Amblyomma* ticks, particularly the males, often display conspicuous ornamentation, and its function and evolutionary significance are interesting biological questions (Schachat et al., 2018).

### Mitochondrial Genome and phylogenetic relationships

We describe the mitochondrial genome of *Amblyomma sparsum* that shows the same arrangement and similar size as the rest of Metastriata (Kelava et al., 2021). Its base composition of 21.1% GC is similar to that of *A. marmoreum* and *A. nuttalli* (Mans et al., 2019; Kelava et al., 2023). The gene order and the two control regions typically found in metastriate ticks but absent in *Africanella transversale* (Kelava et al., 2021) are retained.

This is the first phylogenomic study to show a close relationship between *A. sparsum* and *A. nutalli*. It also confirms the close relationship of *A. tholloni* and *A. hebraeum* with the *A. marmoreum* complex previously reported in mitogenomic (Kelava et al., 2021; Cotes-Perdomo et al., 2023) and other molecular phylogenies (Hornok et al., 2020; Uribe et al., 2020). Unfortunately, apart from *A. marmoreum* itself, there is little mitogenomic data available for other members of the complex and what is available (Kushimo, 2013, Beati and Keirans, 2001) is from different gene segments, making a valid phylogenetic analysis impossible. However, *A. sylvaticum* sequences were found to be highly divergent from our sequence (data not shown) confirming previous reports and suggesting that *A. sylvaticum* may not belong to the *A. marmoreum* complex. As in previous studies it was also not possible to resolve the phylogenetic relationship with member of related complex, such as *A. testudinarium, A. javanense, A. fimbriatum, A. gervaisi* and *A. latum*. More comprehensive and comparable datasets are therefore needed to resolve the taxonomic structure of the *A. marmoreum* complex, its relationship with related complexes and *A. sylvaticum* and the subgeneric status of *Xiphiastor* to which the *A. marmoreum* complex is assigned, but which does not appear to be monophyletic (Uribe et al., 2020; Kelava et al., 2021, 2023). The utilization of robust datasets has proven beneficial for resolving the phylogeny of other *Amblyomma* complexes (Mohamed et al., 2022; Cotes-Perdomo et al., 2023).

### Spotted fever genome of *Rickettsia africae*

We were able to show that the *Amblyomma sparsum* specimen we analysed contained *Rickettsia africae* and we were able to obtain a genome with similar completeness (74.5%) and high similarity (98.7%) to the previously reported *R. africae* strain ESF-5 (CP001612; Fournier et al., 2009). We name this strain AM14. We also obtained a circular plasmid sequence, which contains eight protein-coding genes and has 99.83% similarity to that found associated with *R. africae* ESF-5, but which is shorter, with four fewer coding genes. This strongly suggests that our plasmid belongs to *R. africae* AM14. Phylogenomic analysis places *R. africae* Senegal-AM14 as sister to strain ESF-5 in the Spotted Fever Groups II (SFGII) (El Karkouri et al., 2022), with its next nearest relative being *R. parkeri*.

Metagenomic approach is proving increasingly powerful for detecting microorganisms and retrieving genomic information for evolutionary studies. However, culture-based isolation of pure strains is still necessary for some analyses, such as confirming the association of mobile elements such as plasmid (Hu et al., 2011), and theoretically speaking it is not possible to prove that the plasmid we detected is an episome of *R. africae* AM14 without obtaining a pure isolate. For an accurate understanding of the evolution, geographic origin, and diversification, population dynamics and pathogenicity of different strains a combination of metagenomic and culture-base methods is more powerful than either method alone (Shi et al., 2010; Wang et al., 2014).

*Rickettsia africae* (Zhang et al., 2023) is the etiological agent of African tick-bite fever (ATBF), a neglected emerging disease in rural zones (Mediannikov et al., 2010) and the second most important cause of systemic febrile illness in travelers after malaria (Freedman et al., 2006). An estimated 3.6 billion people are at risk of infection. The principle recognized vectors are the African *Amblyomma* species *A. variegatum* and *A. habraeum* (Petney et al., 1987; Kelly et al., 1996); they are highly efficient transmission vectors and transmission rates over 90% have been reported (Kelly and Mason, 1991; Socolovschi et al., 2009). However, ATFB may no longer be confined to Africa, as surveys conducted in the Americas, Europe, and Oceania have reported *R. africae*-infected ticks, primarily introduced *A. variegatum* (Kelly et al., 2010; Eldin et al., 2011; Andoh et al., 2015; Vogel et al., 2018; Battisi et al., 2020; Pintore et al., 2021). *Rickettsia africae* seems to be well-adapted its arthropod hosts; it shows no negative effects on them, and a symbiotic relationship with hard ticks is possible (Fournier et al., 2009).

*Rickettsia africae*-like DNA has previously been detected in *Amblyomma sparsum* ticks removed from tortoises in Zambia and Kenya (Andoh et al, 2015; Omondi et al, 2017) but as far as we are aware, this is the first record of *R. africae* in *A. sparsum* ticks on a puff adder or in Senegal. Other records of *R. africae* DNA also involve ticks of the *Rhipicephalus, Hyalomma*, and *Haemaphysalis* genera (Mazhetese et al., 2021). *Rickettsia massiliae*, another SFG *Rickettsia*, has been reported in ticks of the *A marmoreum* complex such as *A. sylvaticum* and *A. marmoreum* collected from reptiles (Mofokeng et al., 2022).

It is important to stress that the mere presence of *R. africae* DNA in *A. sparsum* does not prove that it is a competent vector; for this, transmission has to be demonstrated. It is therefore too early to draw any firm conclusions regarding the role of *A. sparsum* in the epidemiology of *R. africae*.

Previous reports of *R. africae* in ticks associated with reptiles, coupled with our findings, suggest that the role of reptiles and *Amblyomma* ticks in the enzootic cycle of ATBF may be underestimated (Sánchez-Montes et al., 2019). However, whether the various reptilian hosts amplify, dilute or kill SFG-rickettsiae remains to be determined (Mendoza-Roldan et al., 2021; Pillay et al., 2022, Lane and Quistad, 1998; Randolph and Dobson, 2012; Ticha et al., 2016). Further research is needed and *A. sparsum* and *B. arietans* should be included in these studies.

Species discrimination within a species complex is often challenging, particularly in immature ticks where morphological characteristics are not fully developed or in poor quality specimens where details of morphological characteristics such as enameling may be lost. Microscopic features such as punctation patterns may help to resolve such difficulties. However, where mitochondrial genomes are available, ticks may be identified by molecular methods such as PCR and DNA sequencing. The availability of comprehensive collections of mitochondrial genomes together with appropriate nuclear gene sequences would also allow resolution of phylogenetic relationships among the *Amblyomma* species and their relationship with the other genera within the Metastriata. In this study we show that metagenomic techniques can provide reliable data for such studies, while simultaneously providing microbiome data of potential epidemiological importance. In addition to its potential relevance to the epidemiology of ATBF the finding of *R. africae* in *Amblyomma sparsum* and other ticks that parasitize reptiles underscores the public health risks associated with the trade of wild fauna. Further studies are needed to evaluate the prevalence of *R. africae* in *A. sparsum*, the vector competence of the tick and its phenology in different populations in order to determine transmission risks and develop appropriate preventive strategies.

## Acknowledgments

All laboratory work was conducted in and with the support of the Molecular Systematics Lab (https://www.mncn.csic.es/es/investigación/servicios-cientifico-tecnicos/laboratorio-de-sistematica-molecular) of MNCN-CSIC. Computing was conducted on the Smithsonian High Performance Cluster (https://doi.org/10.25572/SIHPC). Many thanks to the Comunidad de Madrid for Atracción de Talento contract (REFF 2019-T2/AMB-13166) of JEU; ANID National Doctorate Scholarship Program (REFF 2020–21200182). We are also grateful to Gabriel Martínez del Mármol and Alejandro Carreras for their company during the fieldwork campaigns.

## Supplementary Material Captions

**Supplementary Material 1**. List of mitochondrial genomes employed for the phylogenetic analyses presented in Figure 4, left tree.

**Supplementary Material 2**. List of mitochondrial genomes employed for the phylogenetic analyses presented in Figure 4, left tree.

**Supplementary Material 3**. Summary graphic of the BUSCO assessment using alphaproteobacteria_odb10 (Creation date: 2020-03-06, number of genomes: 744, number of BUSCOs: 432) database.

**Supplementary Material 4**. Gene annotations of the complete mt genomes of *Amblyomma sparsum*.

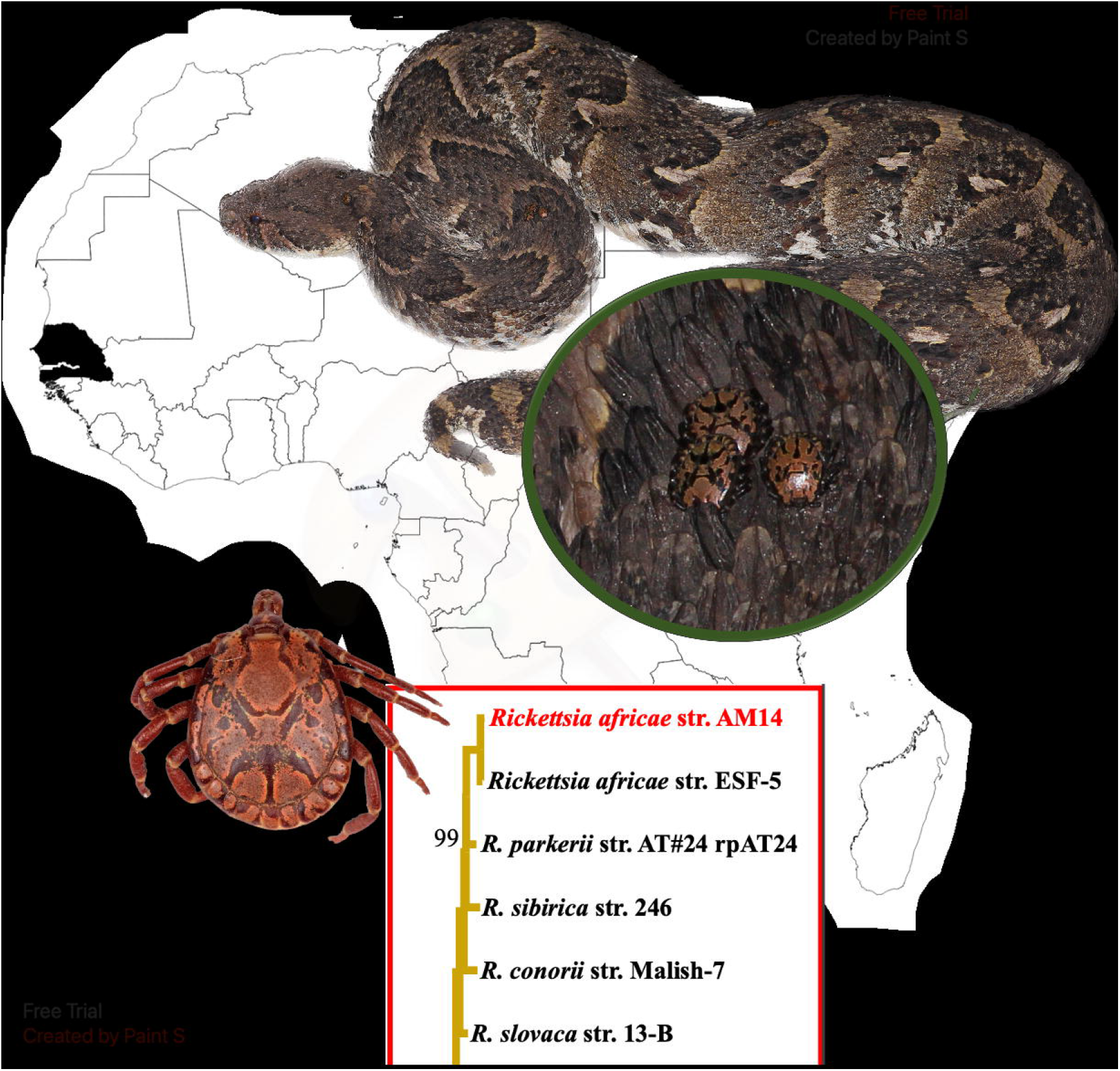

