## Supplementary Material 1-4 for "New insights into the systematics of the Afrotropical *Amblyomma marmoreum* complex (Acari, Ixodidae) and a novel *Rickettsia africae* strain using morphological and metagenomic approaches"

**Supplementary Material 1**. List of mitochondrial genomes employed for the phylogenetic analyses presented in Figure 4, left tree.

| **Spacie** | **Length** | **Acc. Number** | **Reference** |
|---|---|---|---|
| *Amblyomma sparsum* | 14679 | OQ842962 | **This study** |
| *Amblyomma hebraeum* | 14654 | KY457513 | Mans et al. (2019) |
| *Amblyomma marmoreum* | 14676 | KY457515 | Mans et al. (2019) |
| *Amblyomma marmoreum* | 14677 | KY457516 | Mans et al. (2019) |
| *Amblyomma tholloni* | 14642 | KY457521 | Mans et al. (2019) |
| *Hyalomma asiaticum asiaticum* | 14720 | MF101817 | Liu et al. (2018) |
| *Amblyomma geoemydae* | 14780 | MK814531 | Chang et al. (2019) |
| *Amblyomma testudinarium* | 14760 | MT029329 | Chang et al. (2020) |
| *Amblyomma maculatum* | 14803 | MW719251 | Brenner and Raghavan (2021) |
| *Hyalomma rufipes* | 14761 | MW884229 | Lang et al. (2022) |
| *Rhipicephalus sanguineus* | 14710 | NC_002074 | Black and Roehrdanz (1998) |
| *Amblyomma triguttatum* | 14740 | NC_005963 | Shao et al. (2005) |
| *Amblyomma fimbriatum* | 14705 | NC_017759 | Burger et al. (2012) |
| *Rhipicephalus microplus* | 14905 | NC_023335 | Burger et al. (2014) |
| *Dermacentor nitens* | 14839 | NC_023349 | Burger et al. (2014) |
| *Dermacentor silvarum* | 14945 | NC_026552 | Guo et al. (2016) |
| *Amblyomma americanum* | 14709 | NC_027609 | Williams-Newkirk et al. (2015) |
| *Rhipicentor nuttalli* | 14779 | NC_039828 | Mans et al. (2019) |
| *Dermacentor everestianus* | 15191 | NC_042764 | Yu et al. (2018) |
| *Amblyomma javanense* | 14780 | NC_043872 | Duan et al. (2020) |
| *Amblyomma ovale* | 14760 | NC_050255 | Uribe et al. (2020) |
| *Amblyomma gervaisi* | 14709 | OL741734 | Kelava et al. (2023) |
| *Amblyomma latum* | 14658 | OL741735 | Kelava et al. (2023) |
| *Amblyomma nutallli* | 14681 | OL741736 | Kelava et al. (2023) |
| *Amblyomma cajenense* | 14776 | OP901701 | Cotes-Perdomo et al. (2023) |
| *Amblyomma mixtum* | 14815 | OP901702 | Cotes-Perdomo et al. (2023) |
| *Amblyomma patinoi* | 14779 | OP901704 | Cotes-Perdomo et al. (2023) |
| *Amblyomma tonelliae* | 14731 | OP901705 | Cotes-Perdomo et al. (2023) |
| *Amblyommaaa sculptum* | 14784 | OP901706 | Cotes-Perdomo et al. (2023) |

**Supplementary Material 2.** List of mitochondrial genomes employed for the phylogenetic analyses presented in Figure 4, left tree.

| Species | Strain | BioSample | BioProject | Assembly | Size (bp) | GC% | Scaffolds | CDS | Country | Host | Publications |
|---|---|---|---|---|---|---|---|---|---|---|---|
| *Rickettsia aeschlimannii* | | SAMEA2501490 | PRJEB6087 | GCA_001051325 | 1312196 | | 16 | | | | Urmite Genomes, 2014 (unpublished) |
| *Rickettsia africae* | ESF-5 | SAMN02603146 | PRJNA18269 | GCF_000023005 | 1278540 | 33,46 | 1 | 1443 | Ethiopia: Shulu province | *Amblyomma variegatum* | Fournier et al., 2009. Analysis of the Rickettsia africa |
| ***Rickettsia africae*** | **Senegal-M14** | **SUB13049563** | **PRJNA951762** | | **1281475** | **32,4** | **1** | **1434** | **Senegal: Southwest** | ***Amblyomma sparsum*** | **This study** |
| *Rickettsia amblyommatis* | GAT-30V | SAMN02603561 | PRJNA75043 | GCA_000284055 | 1407796 | 32,45 | 4 | 1643 | Stockbridge, GA, USA | *Amblyomma americanum* | Karpathy et al., 2016. Rickettsia amblyommatis sp. n |
| *Rickettsia argasii* | T170-B | SAMN02666743 | PRJNA224116 | GCF_000965185 | 1437875 | 32,30 | 30 | 1429 | USA | *Homo sapiens* | Daugherty et al., 2015 (unpublished) |
| *Rickettsia australis* | Cutlack | SAMN02603558 | PRJNA75037 | GCA_000284155 | 1296670 | 32.33 | 1 | 1402 | | | Johnson et al., 2012 (unpublished) |
| *Rickettsia bellii* | RML Mogi | SAMN02666639 | PRJNA212474 | GCA_000965045 | 1618071 | 31,5 | 4 | 1619 | Brazil | *Amblyomma aureolatum* | Daugherty et al., 2015 (unpublished) |
| *Rickettsia canadensis* | CA410 | SAMN02603555 | PRJNA75029 | GCA_000283915 | 1150228 | 31 | 1 | 1013 | | | Johnson et al., 2012 (unpublished) |
| *Rickettsia conorii* | Malish 7 | SAMN02603141 | PRJNA42 | GCA_000007025 | 1268755 | 32,4 | 1 | 1419 | | | Ogata et al., 2000. Selfish DNA in protein-coding ge |
| *Rickettsia conorii* subsp. *heilongjiangensis* | 054 | SAMN02602971 | PRJNA66907 | GCA_000221205 | 1278471 | 32.30 | 1 | 1424 | China: Heilongjiang province | *Dysmicoccus sylvarum* | Duan et al., 2011. Complete genome sequence of Ric |
| *Rickettsia conorii* subsp. *raoultii* | Khabarovsk | SAMN03372504 | PRJNA276402 | GCA_000940955 | 1344642 | 32.63 | 4 | 1649 | Russia | *Dermacentor silvarum* | El Karkouri et al., 2016. Genome sequence of the tic |
| *Rickettsia conorii* subsp. *raoultii* | IM16 | SAMN06249243 | PRJNA362804 | GCA_001975185 | 1344557 | 32.50 | 1 | 1528 | China: Inner Mongolia | *Homo sapiens* | Cao et al., 2016 (unpublished) |
| *Rickettsia felis* | URRWXCal2 | SAMN02603143 | PRJNA13884 | GCA_000012145 | 1485148 | 33,60 | 1 | 1512 | USA: California | | Ogata et al., 2005. The genome sequence of Rickettsi |
| *Rickettsia fournieri* | AUS118 | SAMEA104430940 | PRJNA224116 | GCF_900243065 | 1447347 | 32,40 | 6 | 1538 | Australia | *Argas lagenoplastis* | Urmite Genomes, 2015 (unpublished) |
| *Rickettsia gravesii* | BWI-1 | SAMN02472051 | PRJNA224116 | GCF_000485845 | 1347499 | 32,20 | 29 | 1538 | Australia | *Amblyomma triguttatum* | Sentausa et al., 2013. Genome sequence of Rickettsia |
| *Rickettsia helvetica* | C9P9 | SAMN02471333 | PRJNA82855 | GCF_000255355 | 1417015 | 32,64 | 2 | 1592 | Switzerland | *Ixodes ricinus* | Dong et al., 2012. Genomic comparison of Rickettsia |
| *Rickettsia honei* | RBT | SAMN02469768 | PRJNA158665 | GCA_000263055 | 1268758 | 32 | 11 | 1462 | Australia: Flinders Island | *Homo sapiens* | Xin et al., 2012. Genomic comparison of Rickettsia h |
| *Rickettsia japonica* | YH | SAMD00060967 | PRJDA38487 | GCA_000283595 | 1283087 | 32.40 | 1 | 1448 | Japan | *Homo sapiens* | Matsutani et al., 2013. Complete genomic DNA seque |
| *Rickettsia massiliae* | MTU5 | SAMN02603147 | PRJNA18271 | GCA_000016625 | 1360898 | 32,49 | 2 | 1544 | | | Blanc et al., 2007. Lateral gene transfer between obli |
| *Rickettsia monacensis* | IrR/Munich | SAMEA4532341 | PRJNA224116 | GCA_000499665 | | 32.4 | 1 | 1519 | England | | Felsheim et al., 2013 (unpublished) |
| *Rickettsia montanensis* | OSU 85-930 | SAMN02603562 | PRJNA75045 | GCF_000284175 | 1279798 | 32,6 | 1 | 1413 | | | Johnson et al., 2012 (unpublished) |
| *Rickettsia parkeri* | AT#24 | SAMN02666644 | PRJNA212479 | GCA_000965075 | 1300534 | 32,4 | 1 | 1438 | Brazil | *Amblyomma triste* | Daugherty et al., 2015 (unpublished) |
| *Rickettsia peacockii* | Rustic | SAMN02604073 | PRJNA31309 | GCF_000021525 | 1288492 | 34,66 | 1 | 1456 | USA | | Felsheim et al., 2009. Genome sequence of the endos |
| *Rickettsia philipii* | 364D | SAMN02603554 | PRJNA75027 | GCA_000283995 | 1287740 | 32,5 | 1 | 1414 | | | Johnson et al., 2012 (unpublished) |
| *Rickettsia prowazekii* | RpGvF24 | SAMN02603547 | PRJNA75009 | GCA_000277265 | 1112101 | 29 | 1 | 850 | | | Johnson et al., 2012 (unpublished) |
| *Rickettsia rhipicephali* | HJ#5 | SAMN04240637 | PRJNA301161 | GCA_001442475 | 1406075 | 32.26 | 3 | 1592 | Brazil: Atlantic rain forest , Sao Paulo | *Haemaphysalis juxtakochi* | Felsheim et al., 2015 (unpublished) |
| *Rickettsia rhipicephali* | 3-7-female6-CWPP | SAMN02603560 | PRJNA75041 | GCA_000284075 | 1290368 | 32.39 | 2 | 1452 | | | Johnson et al., 2012 (unpublished) |
| *Rickettsia rhipicephali* | Ect | SAMN02585079 | PRJNA212462 | GCA_000964905 | 1266919 | 32.60 | 1 | 1421 | USA | *Rhipicephalus sanguineus* | Daugherty et al., 2015 (unpublished) |
| *Rickettsia rickettsii* | Morgan | SAMN03324096 | PRJNA205204 | GCA_000831545 | 1269809 | 32,5 | 1 | 1405 | USA: Kannapolis | *Homo sapiens* | Clark et al., 2015. Comparative genome sequencing |
| *Rickettsia sibirica* | 246 | SAMN02393728 | PRJNA1414 | GCA_000166935 | 1250021 | 32,5 | 1 | 1392 | Russia: Krasnojarsk | | Malek et al., 2004. Protein interaction mapping on a |
| *Rickettsia slovaca* | 13-B | SAMN02603145 | PRJNA15712 | GCA_000237845 | 1275089 | 32,5 | 1 | 1444 | Europe | *Dermacentor marginatus, Dermacentor reticulatus* | Fournier et al., 2012. Complete genome sequence of |
| *Rickettsia* sp. | Humboldt | SAMN02665668 | PRJNA232537 | GCA_000965155 | 1482156 | | 1 | 1739 | USA | Ixodes pacificus | Daugherty et al., 2015 (unpublished) |
| *Rickettsia typhi* | B9991CWPP | SAMN02603530 | PRJNA65221 | GCA_000277305 | 1112957 | 28,9 | 1 | 813 | Myanmar | *Bandicota* sp. | Elliott, Ivo, et al. "Oxford nanopore MinION sequen |

**Supplementary Material 3.** Summary graphic of the BUSCO assessment using alphaproteobacteria_odb10 (Creation date: 2020-03-06, number of genomes: 744, number of BUSCOs: 432) database.

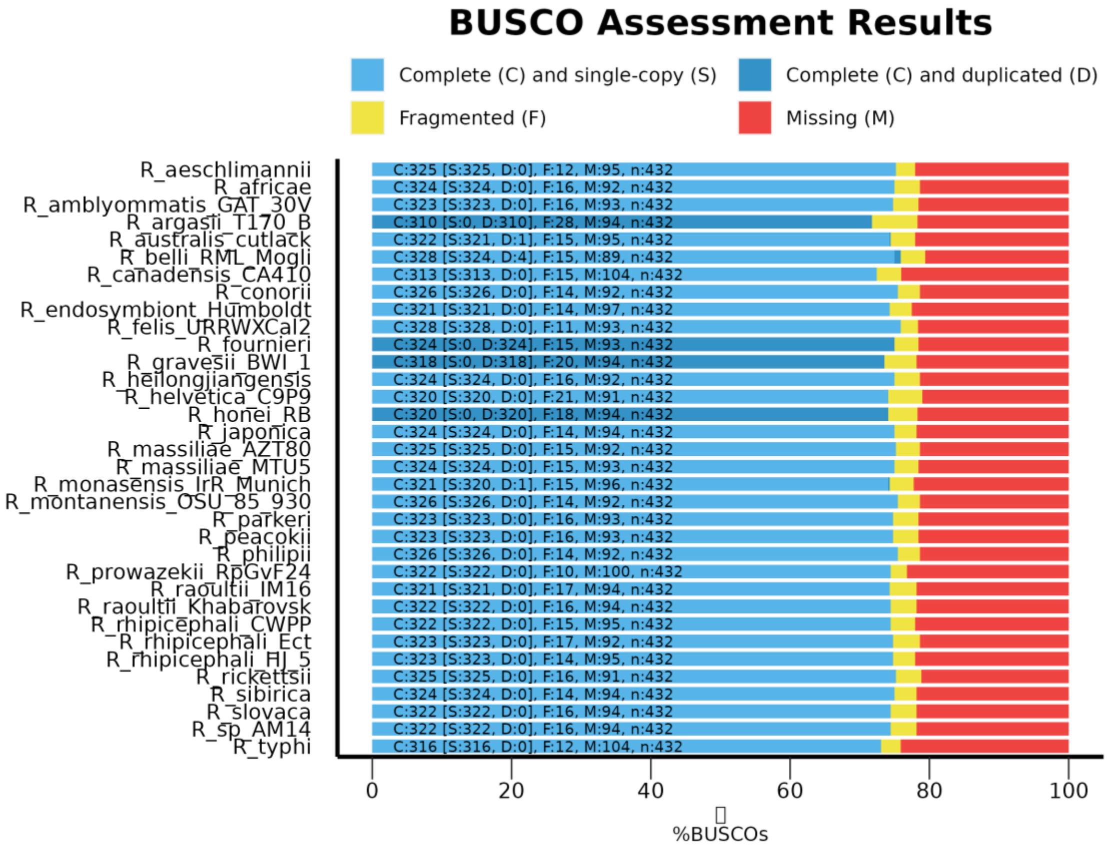

**Supplementary Material 4.** Gene annotations of the complete mt genomes of *Amblyomma sparsum*.

| Type | Gene | Start | Stop | Length | Stop codon | Strain |
|---|---|---|---|---|---|---|
| tRNA | Met | 1 | 62 | 62 | | forward |
| CDS | ND2 | 63 | 1,016 | 954 | TAA | forward |
| tRNA | Trp | 1,015 | 1,076 | 62 | | forward |
| tRNA | Tyr | 1,075 | 1,136 | 62 | | reverse |
| CDS | COX1 | 1,144 | 2,667 | 1,524 | TAA | forward |
| CDS | COX2 | 2,674 | 3,348 | 675 | TAA | forward |
| tRNA | Lys | 3,349 | 3,414 | 66 | | forward |
| tRNA | Asp | 3,414 | 3,477 | 64 | | forward |
| CDS | ATP8 | 3,478 | 3,636 | 159 | TAA | forward |
| CDS | ATP6 | 3,63 | 4,292 | 663 | TAA | forward |
| CDS | COX3 | 4,296 | 5,073 | 778 | T— | forward |
| tRNA | Gly | 5,074 | 5,135 | 62 | | forward |
| CDS | ND3 | 5,136 | 5,477 | 342 | TAA | forward |
| tRNA | Ala | 5,477 | 5,538 | 62 | | forward |
| tRNA | Arg | 5,544 | 5,602 | 59 | | forward |
| tRNA | Asn | 5,602 | 5,671 | 70 | | forward |
| tRNA | Ser | 5,671 | 5,725 | 55 | | forward |
| tRNA | Glu | 5,73 | 5,794 | 65 | | forward |
| CDS | ND1 | 5,788 | 6,726 | 939 | TAA | reverse |
| tRNA | Leu | 6,727 | 6,791 | 65 | | reverse |
| rRNA | 16S rRNA | 6,792 | 7,993 | 1,202 | | reverse |
| tRNA | Val | 7,994 | 8,055 | 62 | | reverse |
| rRNA | 12S rRNA | 8,056 | 8,752 | 697 | | reverse |
| D-loop | D-loop | 8,753 | 9,058 | 306 | | forward |
| tRNA | Ile | 9,059 | 9,123 | 65 | | forward |
| tRNA | Trp | 9,128 | 9,194 | 67 | | reverse |
| tRNA | Phe | 9,198 | 9,256 | 59 | | reverse |
| CDS | ND5 | 9,263 | 10,918 | 1,656 | TAA | reverse |
| tRNA | His | 10,919 | 10,981 | 63 | | reverse |
| CDS | ND4 | 10,981 | 12,306 | 1,326 | TAG | reverse |
| CDS | ND4L | 12,3 | 12,575 | 276 | TAA | reverse |
| tRNA | Thr | 12,578 | 12,638 | 61 | | forward |
| tRNA | Pro | 12,643 | 12,707 | 65 | | reverse |
| CDS | ND6 | 12,714 | 13,142 | 429 | TAA | forward |
| CDS | CYTB | 13,146 | 14,222 | 1,077 | TAG | forward |
| tRNA | Ser | 14,222 | 14,284 | 63 | | forward |
| tRNA | Leu | 14,284 | 14,348 | 65 | | reverse |
| D-loop | D-loop | 14,349 | 14,618 | 270 | | forward |
| tRNA | Cys | 14,619 | 14,672 | 54 | | forward |
